# The cloacal outgrowth orchestrates co-development of the bladder and umbilical arteries

**DOI:** 10.1101/2025.11.18.689093

**Authors:** Xing Ye, Liguang Xia, Xianfa Yang, Ruirong Tan, Haochuan Zhang, Liming Lei, Jungang Huang, Yingsheng Zhang, Edward Morrisey, Ping Zhu, Zhongrong Li, Naihe Jing, Xue Li

## Abstract

The urinary bladder is an innovation of eutherian mammals to manage liquid wastes. However, bladder development and its’ embryonic function remain poorly understood. Here, we report that in both humans and mice the bladder emerges from the cloaca as the rostral outgrowth, which is flanked by the umbilical arteries. Arrest of the outgrowth, as seen in *Shh-*null mouse embryos, causes bladder agenesis and unexpected severe defect of the umbilical arteries, suggesting that the outgrowth epithelial signal *Shh* coordinates development of both the bladder and umbilical arteries. *Wnt2* signaling molecule is restricted to the rostral outgrowth mesenchyme; and its’ expression is dramatically down-regulated in *Shh-*null mutants. Furthermore, *Wnt2-*null mutants develop small bladder and single umbilical artery defects. Findings here uncover an unexpected co-development mechanism of the bladder and umbilical arteries orchestrated by the cloacal outgrowth signals. The co-development paradigm offers a conceptual framework to understand evolution and development of the bladder.

**Significance Statement:** The urinary bladder is found in the eutherian mammals but largely absent from other vertebrates. Innovation of the bladder has been viewed as a necessity to manage liquid metabolic wastes on dry land. In this study, we report that the bladder emerges from the rostral outgrowth of the cloaca. We also report an unexpected developmental link between the bladder and umbilical arteries and, demonstrate that their co-development is orchestrated by the outgrowth epithelial signal *Shh* and mesenchymal signal *Wnt2*. Findings here highlight a possibility that, in addition to managing metabolic wastes, normal bladder development is required for growth of mammalian embryos by supporting development of the umbilical arteries.

## Introduction

The urinary bladder is one of the evolutionary innovations of amniotes: it is found in placental mammals but largely absent from reptiles and birds (1, 2). Having the bladder offers selective advantages in water preservation (1) and predator evasion (3) for terrestrial vertebrates to thrive on dry land. While the bladder is known for managing liquid metabolic wastes afterbirth, formation and function of the bladder before birth are often overlooked and poorly understood.

The role of embryonic bladder, particularly at the stages prior to the renal function remain largely unknown. Mouse bladder emerges several days before the kidneys begin to produce urine (4-8), implying that waste management is unlikely the major function of the embryonic bladder. Similarly, human bladder also emerges prior to the renal function. Indeed, mammalian embryos dispose their metabolic wastes primarily through the umbilical arteries to the chorioallantoic placenta or placenta, where the liquid metabolic wastes are removed maternally. Human birth defects with isolated bladder agenesis or absence of the bladder are rare with the estimate incidence of 1:600,000 (9, 10). Majorities of the affected patients exhibit varying degrees of genital abnormalities, from ambiguous to normal genitalia. Unexpectedly, umbilical artery defects are also observed among patients with bladder agenesis, including single umbilical artery phenotype, absence of a common iliac artery, and iliac artery anomaly (11, 12). Absence of the bladder is often found in sirenomelia birth defects (1.5-4.2 cases per 100,000) and caudal dysgenesis (1-2.5 cases per 100,000) (13, 14). This is consistent with the notion that the cloaca functions as an organizing center to instruct development of the caudal structures, including the bladder, genital tubercle, and the hindlimbs (15-17). Bladder agenesis is also observed in rodents when pregnant females are exposed to Adriamycin or doxorubicin, a chemotherapy medication (18). In this teratogenesis model, the cranial portion of the bladder to the point of entries of Wolffian ducts (WD) is absent (19), suggesting that the main body of the urinary bladder, *i.e.,* the cranial portion to the points of entry of the WD, originates from a unique population of progenitor cells that are particularly sensitive to teratogen exposure. Therefore, the bladder is an integral component of the caudally structures that also include the genital tubercle, hindlimb, and possibly the umbilical arteries and; furthermore, the cloaca is central to normal development of the bladder and is required for mammalian embryos to thrive inside the womb.

The cloaca is a caudal extension and local expansion of the hindgut, which is formed around embryonic day 10.5 (E10.5) in mice and the 4^th^ weeks of gestation in humans (6, 7, 20, 21). For reptiles and birds, which reproduce inside the cleidoic eggs, the embryonic wastes are drained from the cloaca to the allantois, an extraembryonic sac functioning like an embryonic bladder. The cloaca persists after hatching and the adult cloaca serves as the opening shared by the digestive, urinary, and reproductive tracts of reptiles and birds. The cloaca-allantois conduit is conserved among vertebrates. However, unlike the cloaca from reptiles and birds, mammalian cloaca undergoes dramatic morphogenetic changes, in part due to the region-specific recruitment and asymmetric growth of peri-cloacal mesenchyme (PCM) such as intra-cloacal mesenchyme (ICM) or the urorectal septum (URS) as well as the dorsal PCM (dPCM) (22-24). This morphologic transformation divides the cloaca into two halves: the ventral half or urogenital sinus (UGS) differentiates into the lower urinary tract while the dorsal half contributes to the digestive tract (6, 7, 20). Consequently, the urinary and digestive tracts are connected in reptiles and birds but separated in mammals. For reasons that are largely unknown, the bladder emerges at the junction between the UGS and the allantois in placental mammals (2, 25, 26).

In a mouse line deficient of transmembrane protein 132a (*Tmem132a^-/-^*) (27), the gut fails to extend caudally to form the cloaca at embryonic day 9.5 (E9.5). Consequently, the bladder as well as the genital tubercle are absent from *Tmem132a^-/-^* embryos. Sonic hedgehog (*Shh*) encodes a signaling molecule that is expressed in the cloacal epithelium (28-30). The cloaca is formed initially in *Shh^-/-^* embryos, but the PCM are underdeveloped (6), resulting in persistent cloaca and the hypoplastic bladder and genital tubercle phenotype (28-30). Compared to *Shh^-/-^*mutation alone, compound mutants deficient of both *Shh* and bone morphogenic protein *7* (*Bmp7*) exhibit an even more severe phenotype that resembles human birth defect sirenomelia – the genital tubercle is absent and the hindlimbs are fused (31). The cloaca of *Shh;Bmp7* compound mutants is narrower than normal and shows a reduced local expansion at E9.5 (31). Sirenomelia phenotype is also observed in two other mouse models including a compound mouse mutant line lacking both *Bmp7* and twisted gastrulation (*Tsg*) (32) and a mouse line with conditional knockout of *Bmp4* (33). Therefore, formation of the cloaca is the critical first step towards formation of the bladder. How the bladder emerges from the cloaca or, specifically, the junction between the UGS and allantois, remains largely unknown.

Here, we investigate bladder development in humans and mice. Our study uncovers an unexpected co-development mechanism of the bladder and umbilical arteries, which is controlled in part by the cloacal outgrowth epithelial and mesenchymal signals. The co-development paradigm offers a conceptual framework to understand development and evolution of mammalian caudal structures as well as human birth defects such as bladder agenesis and sirenomelia.

## Results

### Mammalian bladder originates from the cloaca instead of the allantois

It has been postulated that the urinary bladder originates from both the cloaca and the allantois, with the neck or proximal part of the urinary bladder deriving from the cloaca while the remainder of the bladder, such as the bladder dome, from the allantois (2, 26). To test this hypothesis directly, we tracked developmental trajectory of the allantois of human and mouse embryos using a comprehensive series of digitized high-resolution episcopic microscope (HREM) three-dimension (3D) images (Table S1 and S2) (6-8). In human embryos, formation of the cloaca-allantois canal was observed initially at Carnegie Stage 8 (CS8) as an endoderm protrusion that extended into the body stalk (Figures S1A-D). The canal was clearly visible at CS12 (Figure 1A-D), which connects the cloaca directly to the allantois to form a continuous embryonic-extraembryonic conduit. Unlike what are observed in bird and reptile embryos (25), however, human allantoic gut was only a rudimentary structure and became progressively smaller at later stages at CS14, 18, and 21 (Figure S2, video).

**Figure 1.**
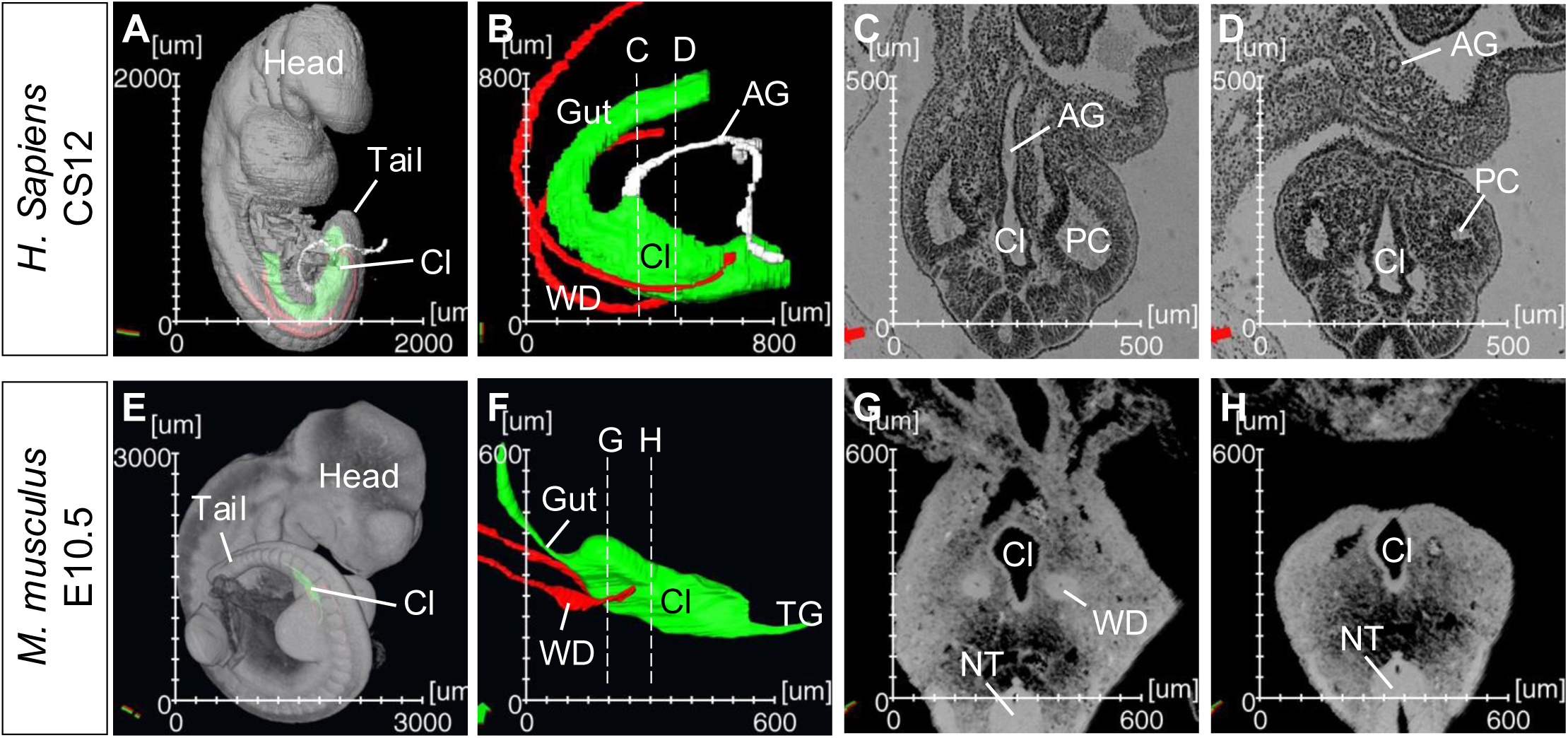
Species variations of the allantoic gut. The allantoic gut (AG) is found in human (A-D) but not mouse (E-H) allantois. **A** and **E**, Wholemount ventrolateral views of CS12 human (A) and E10.5 mouse (E) embryos; **B** and **F**, 3D segmentation of the definitive gut and cloacal epithelium (green), allantoic gut (white), and Wolffian Duct (red); **C, D, G,** and **H,** Cross-sections of the cloaca taken from planes shown in B and F (dash lines). AG, Allantoic gut; Cl, Cloaca; NT, Neural Tube; PC, Peritoneal Cavity; TG, Tail Gut; WD, Wolffian Duct.

Surprisingly, the cloaca-allantois canal was not observed in mouse embryos at the comparable stages (Figures 1E-H, and Figure S3 video). Instead, mouse cloaca ended blindly at the caudal end of the trunk. To rule out a possibility that the presumptive allantoic gut might have formed initially, but was degenerated thereafter, we examined E8.5 mouse embryos, a stage comparable to CS8 human embryos. Unlike human embryos, an expected endodermal protrusion was never observed in mouse embryos based on the HREM images (Figures S1E-H). To ascertain this finding, we analyzed serial histological sections, which showed that the hindgut was formed, but ended blindly ventral to the caudal neuropore of E8.5 mouse embryos (Figures S1I-K). In comparison, the allantois at this stage (E8.5) had already established an elaborate extraembryonic vascular network with no evidence of the cloaca-allantois canal. Absence of the cloaca-allantois canal in mouse embryos raises a strong possibility that the entire urinary bladder originates from the cloaca.

### The bladder emerges as the rostral outgrowth of the cloaca

To further examine the possibility, we asked how the bladder may emerge from the cloaca in human and mouse embryos. First, we used the Wolffian ducts (WD) insertion sites as the spatial references because they have been considered as the stable anatomic landmarks along the rostrocaudal axis of the cloaca (7). In mouse embryos, the WDs inserted into the cloaca at E10.5 (Figure 2A). The UGS is the ventral half of the cloaca (E11.5 and E12.5) oriented along the rostro-caudal axis (Figures 2B-D). At E12.5, the rostral outgrowth of the cloaca was observed, which inserted into the triangular mesenchymal space (Figure 2C, insert). By E13.5, the rostral outgrowth became a prominent hollow structure, an appearance of the bladder as a distinct anatomical structure (Figure 2D). Concomitant to formation of the bladder at the rostral end of the cloaca, a caudal extension of the cloaca was also observed, resulting in formation of the genital tubercle (Figures 2C and D).

**Figure 2.**
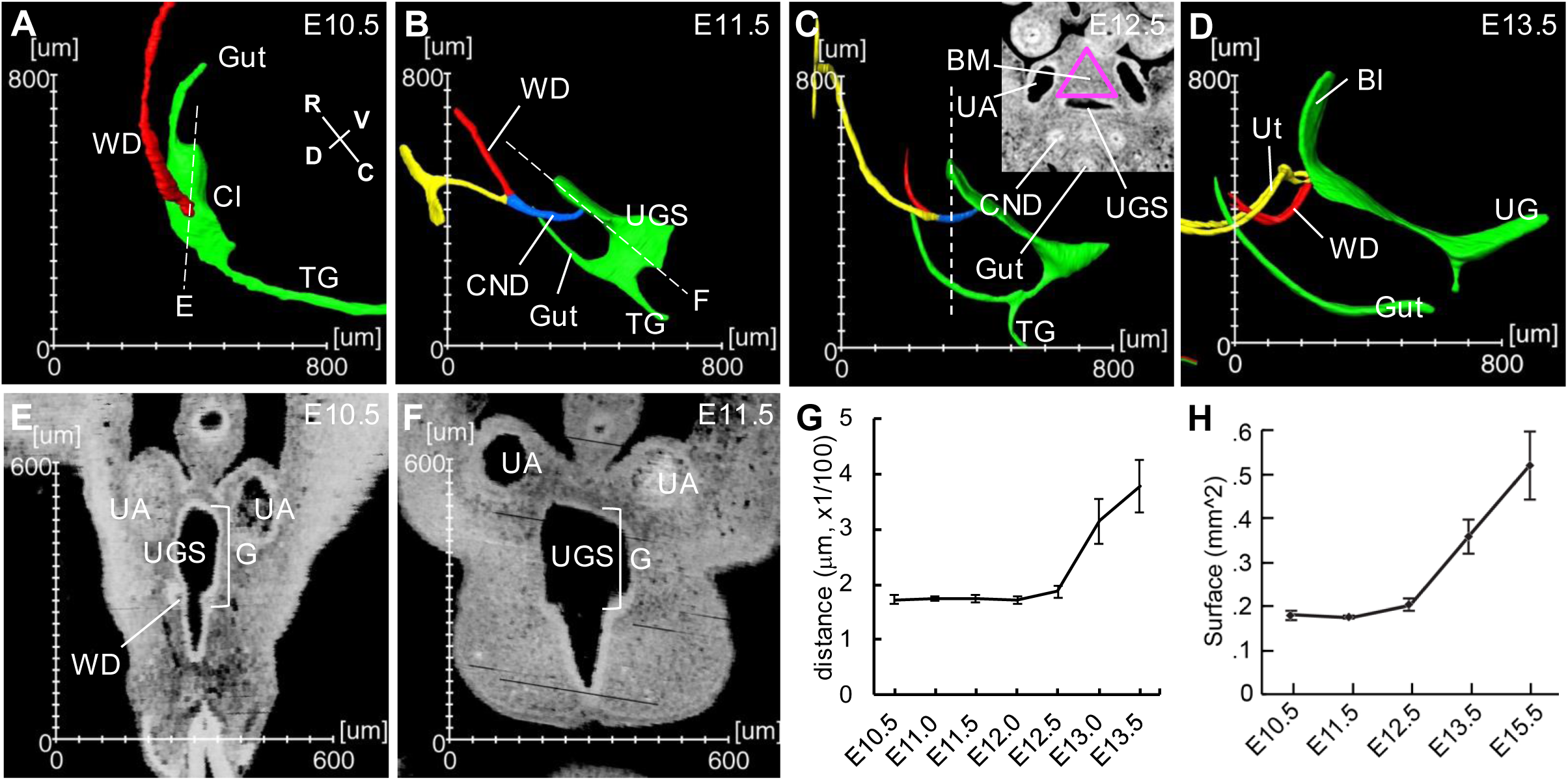
Rostral outgrowth of the cloaca forms the bladder. **A-D**, 3D sagittal views of mouse cloaca from E10.5 to E13.5. Insert in C, a digital cross section from the plane shown as dashed lines. The triangular mesenchymal domain is marked in purple. **E** and **F**, Digital sections through the planes shown in A and B, respectively. Bl, Bladder; CND, common nephric duct; UA, umbilical artery; UGS, urogenital sinus; Ut, ureter; and more abbreviations in Figure 1. **G**, Distance from insertion site of Wolffian Duct to the rostral tip of the cloaca as shown in E and F. **H**, Surface area of the cloaca.

These observations suggest that the bladder emerges from the rostral outgrowth of the cloaca. To substantiate this conclusion, we measured the distance from the WD insertion sites to the tip of rostral outgrowth and the epithelial surface area of the entire cloaca (Figures 2E-H). The distance remained constant from E10.5 (∼172.67 μm) to E12.5 (∼187.33 μm) but increased substantially from E12.5 (187.33 μm) to E13.0 (∼314.67 μm) to E13.5 (∼378.33 μm) (Figure 2G). The overall surface area of the cloaca was unchanged from E10.5 to E12.5 but began to increase at E12.5 (Figure 2H). These findings suggest that transformation of the cloaca proceeds the outgrowth, and further confirm that the outgrowth of the cloaca proceeds formation of the bladder rostrally and genital tubercle caudally.

The umbilical arteries have been used as another independent anatomic landmarks along the rostrocaudal axis (7, 34). The human cloaca-allantois canal traversed between the umbilical arteries (Figures 3A-D). The junction between the cloaca and allantois, which is marked by the transition from pseudostratified columnar epithelium to low cuboidal epithelium, respectively (7), was located near the root of umbilical arteries (Figures 3A-D, arrows). A rostral expansion of the cloaca was visible at CS18 and progressed to form the bladder flanked by the umbilical arteries (Figures 3C and D). In mouse embryos, the rostral outgrowth was also flanked by the umbilical arteries at the comparable developmental stages (Figure 3E-H). The rostral tip of the cloaca located near the umbilical arteries at E10.5 and E11.5. By E13.5 and 15.5, the bladder was formed between the umbilical arteries. Together, these findings support the notion that the bladder emerges from the rostral outgrowth of the cloaca in humans and mice.

**Figure 3.**
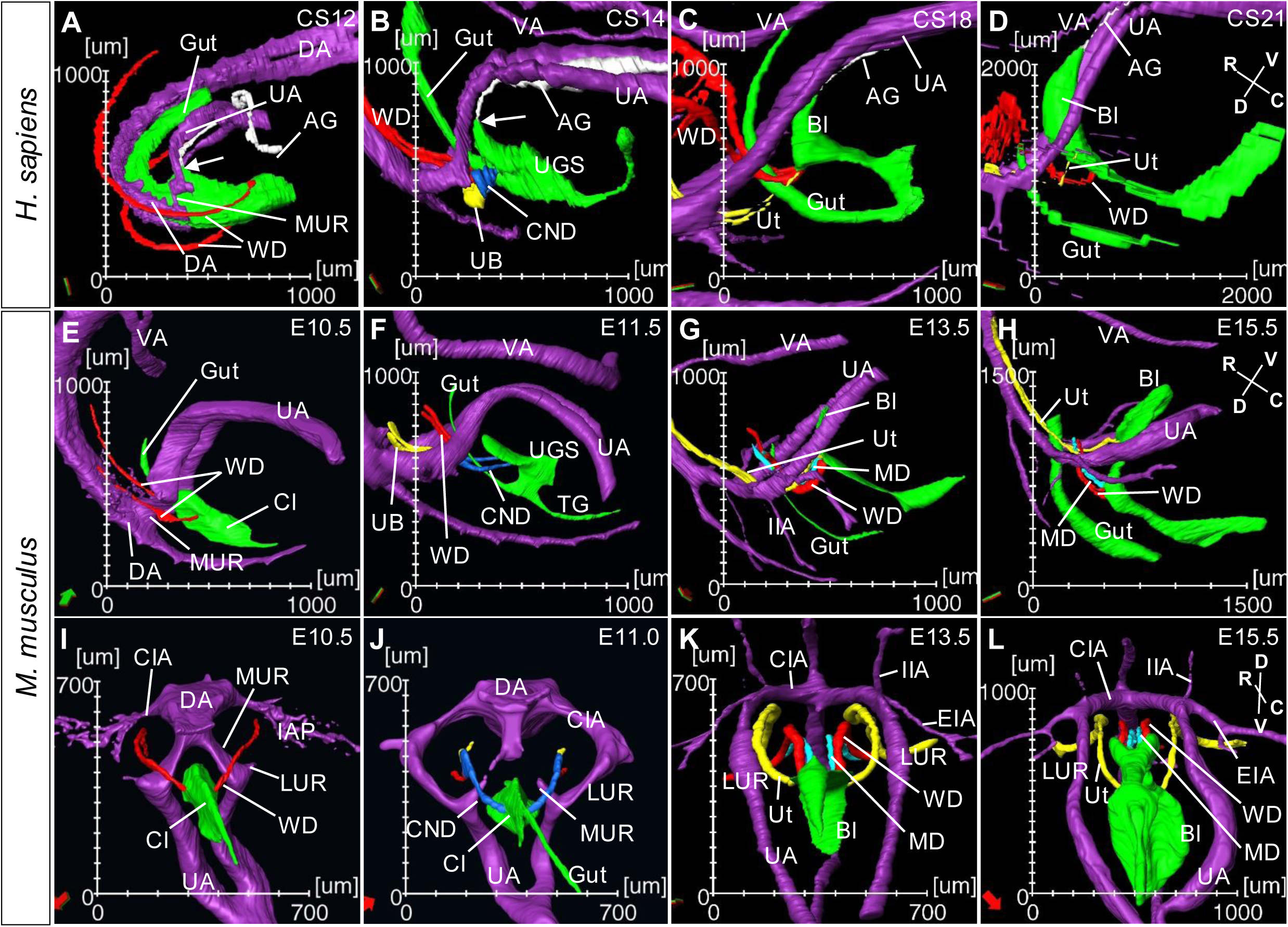
Spatiotemporal correlation of the rostral outgrowth of the cloaca and remodeling of the umbilical arteries in human and mouse. **A-H**, The 3D sagittal views of outgrowth of the cloaca and remodeling of the umbilical arteries in human (A-D) and mouse (E-H) embryos at the comparable developmental stages. **I**-**L**, Caudal views of mouse umbilical arteries reveal the critical periods of umbilical artery remodeling between E10 and E11, and maturation of the iliac arteries between E13-E15 in mice. CIA, common iliac artery; DA, dorsal aorta; EIA, external iliac artery; IIA, internal iliac artery; LUR, lateral umbilical artery root; MUR, medial umbilical artery root; UB, ureteric bud; VA, vitelline artery; more abbreviations in Figures 1 and 2.

### The rostral outgrowth links development of the bladder and umbilical arteries

The umbilical arteries are the vital structures that carry deoxygenated blood and metabolic wastes from the fetus to the placenta. Physical proximity between the developing bladder and umbilical arteries prompted us to examine whether these two seemingly unrelated structures are developmentally linked. During the critical period of the bladder development, the medial umbilical artery roots (MURs) were connected directly to the descending aorta at E10.5 in mice and CS12 in humans (Figures 3A E and I). At this stage, the WDs traversed laterally to the MURs. Surprisingly, the spatial relationship between the WDs and umbilical arteries was reversed at later stages after E11.5, with the WDs traversing medially instead (Figures 3B-D and F-H), indicating a dramatic remodeling of the umbilical arteries.

Possible remodeling of the umbilical arteries became self-evident when embryos were visualized caudally (Figures 3I-L). The MURs were prominent at E10.5 but were degenerated half a day later at E11.0 (comparing Figures 3I and J). The lateral umbilical artery roots (LURs) emerged from the umbilical arteries and the iliac artery plexus (IAP) appeared from the descending aorta at E10.5 (Figure 3I). Concomitant with degeneration of the MURs at E11.0, the mature LURs were formed, which connect the descending aorta indirectly via the comment iliac arteries (CIA) (Figures 3K and L). As the result, the umbilical arteries traversed laterally to the WDs, marking the completion of the umbilical artery remodeling, *i.e.,* degeneration of the MURs and emergence of the LURs and CIA. While mechanism of the umbilical artery remodeling remains to be elucidated, the process of remodeling appears to be conserved in humans and mice.

### *Shh* orchestrates co-development of the bladder and umbilical arteries

To test the hypothesis that the bladder and umbilical arteries are mechanistically linked during embryonic development, we focused on signaling molecule *Shh*, which is restrictively expressed in the cloacal epithelium and absent from the cloacal mesenchyme (35). The cloaca was formed in *Shh* null-mutant (*Shh^-/-^*) embryos at E10.5 (Figure S4, video). The 3D HREM analyses of *Shh^-/-^* embryos also revealed that septation of the cloaca was initiated but the process failed to complete in the mutants, resulting in a persistent cloaca-like phenotype at E13.5. The rostral and caudal outgrowth of the UGS was not observed at E12.5 and, consequently, the urinary bladder and genital tubercle were absent, consistent with the notion that epithelial *Shh* signal is critically required for formation of the bladder and genital tubercle (28-30).

The role of *Hedgehog* signaling in vascular development such as remodeling of the pharyngeal arch arteries and the fetal-placental arterial connection has also been implicated previously (36-38). However, a possible role for *Shh* in umbilical artery development has not been reported. To ascertain that *Shh* is absent from any progenitor cells that may contribute directly to the umbilical arteries, we used the Cre/loxP genetic strategy to indelibly label the *Shh-*expression cells as well as their progenies (39). The labeled cells were visualized whole mount using 3D HREM. Strong signals were detected in all tissues that are known to be the *Shh* genetic lineage, including the gut epithelium, cloacal epithelium, notochord, floorplate, and the zone of polarized activity (Figure S5). Notably, the umbilical arteries were not labeled, indicating that *Shh-*expression cells do not directly contribute to the umbilical arteries.

We next examined whether *Shh* may indirectly affect development of the umbilical arteries in a paracrine fashion. We analyzed a series of *Shh^-/-^* embryos from E10.5 to E13.5 (Figures 4 and S4). The vascular network including the umbilical arteries and dorsal aorta was highlighted to reveal vascular anomalies of *Shh^-/-^*mutants using wild type littermates as controls. The umbilical arteries were observed in *Shh^-/-^*mutants; however, these arteries at E10.5 were apparently smaller in size compared to controls (Figures 4A and E). The IAP was clearly underdeveloped at this stage. Moreover, it failed to progress to form the LURs at later stages between E11.5 and E13.5 (Figures 4B-D and F-H). Unlike wild type controls in which the MURs were degenerated at E11.5, the MURs persisted in *Shh^-/-^* mutants, resulting in the single umbilical artery phenotype at E13.5 (Figures 4, comparing B-D to F-H). The single artery traversed medially to the WD, consistent with a possibility that it originated from the MUR instead of LUR arteries (Figure 4H). Collectively, these findings demonstrate that Shh is a paracrine epithelial signal orchestrating co-development of the bladder and umbilical arteries.

**Figure 4.**
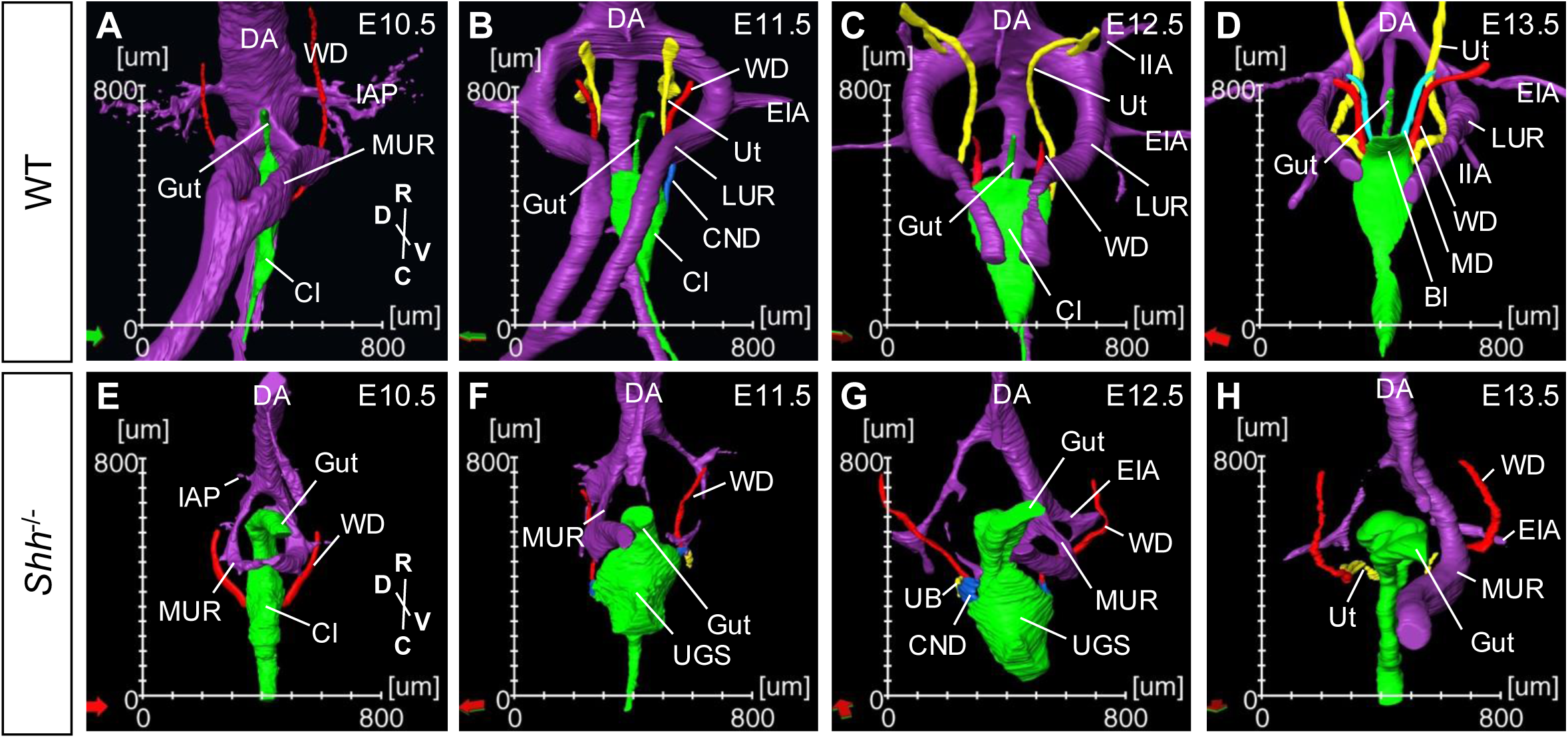
*Shh* is required for remodeling of the umbilical arteries. **A-D**, 3D ventral views of the arterial networks (purple), the cloaca/gut epithelium (green) and associated structures of wild type (A-D) and *Shh^-/-^* mutant (E-H) embryos from E10.5-E13.5. Apparent umbilical artery defects provide evidence that the medial umbilical arteries are lost at one side and the lateral umbilical arteries do not form (F-H). See Figure 3 for abbreviations.

### *Wnt2* is an outgrowth mesenchymal signal required for co-development of the bladder and the umbilical arteries

In addition to the epithelial signal Shh, we hypothesized that co-development of the bladder and umbilical arteries may also depend on mesenchymal signal(s). To test this hypothesis, we performed a spatial transcriptomic analysis of the cloaca (manuscript in preparation). *Wnt2* is among the top candidates, which encodes a canonical Wnt-family signaling molecule and has been independently identified from E16.5 bladder mesenchyme (40). Previous studies have shown that *Wnt2* is expressed in the foregut mesenchyme and regulates lung development (41, 42). However, its potential role in bladder and umbilical artery development has not been explored. We first validated *Wnt2* expression in the outgrowth mesenchyme from E11.5 to E14.5 using wholemount RNA *in situ* hybridization (Figures 5, 6, and S6). In addition to the developing lung, strong *Wnt2* signal was detected in the rostral domain of the cloaca at E11.5 and the developing bladder at E12.5 (Figures 5A and B). The bladder-specific expression pattern became apparent when the stained embryos were examined as histological sections at E13.5 and E14.5 (Figure S6). A sharp *Wnt2-*expression boundary was observed that separated the developing bladder from the developing urethra and genital tubercle (Figures 5B, 6G and S6B), implying that the bladder mesenchyme is molecularly distinctive from the urethral mesenchyme. Together, these findings show that *Wnt2* is highly restricted to the rostral outgrowth mesenchyme, and its expression is maintained in the bladder mesenchyme.

**Figure 5.**
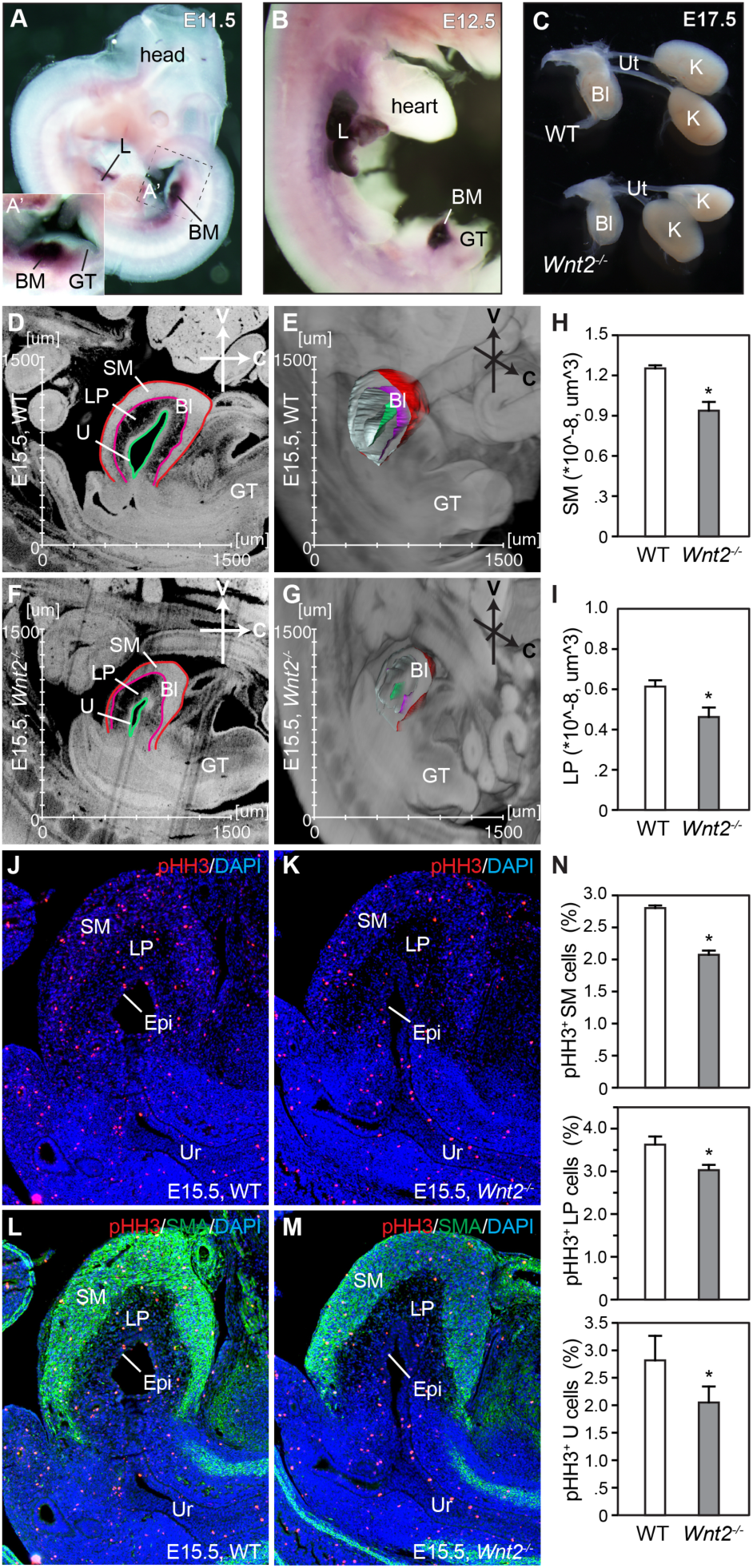
*Wnt2* regulates growth of the bladder. **A** and **B,** *Wnt2* is restrictedly expressed in the developing bladder and lung. Whole mount RNA *in situ* hybridization of mouse embryos at E11.5 (A) and E12.5 (B) using *Wnt2-*specific probe (dark purple staining). **C**, *Wnt2* deficient (*Wnt2^-/-^*) bladders are smaller than controls at E17.5. Whole mount views of the urinary tract. **D**-**I**, HREM images (D-G) and volumetric quantifications (H and I) of the bladders at E15.5 before urine production. Bl, bladder; GT, genital tubercle; LP, lamina propria layer; pBl, primordial SM, smooth muscle layer; U, urothelium. Student’s *t* test, *p<*0.05 (n=4).

*Wnt2* and *Wnt2b* are paralogous genes that share redundant functions in lung development (41, 42). *Wnt2b* is also expressed in the bladder mesenchyme, but at a lower level than *Wnt2* (43). To investigate the potential role of *Wnt2/2b* signaling in co-development of the bladder and umbilical arteries, we examined *Wnt2-*null (*Wnt2^-/-^*) mouse mutants (41, 42). Gross analysis of *Wnt2^-/-^* embryos indicated that the bladder was smaller in size compared to littermate controls (Figure 5C), suggesting the function of *Wnt2* is not fully compensated by *Wnt2b in vivo.* To quantify bladder size, we collected HREM 3D data of E15.5 embryos (Figures 5D-G), one day prior to the urine production. Volumetric analysis of both bladder smooth muscle and lamina propria layers showed a significant reduction in *Wnt2^-/-^*mutants compared to controls (Figures 5H and I). Consistently, cell proliferation rates as indicated by the percentage of pHH3-positive cells were significantly lower in *Wnt2^-/-^* mutants than in controls (Figures 5J-N). The three-dimensional morphometric analysis also revealed that *Wnt2^-/-^* mutants displayed a single umbilical artery phenotype (Figures 6A-D). Unlike wild type controls which had two umbilical arteries traversing bilaterally to the bladder, there was only one umbilical artery found in *Wnt2^-/-^* mutants. Moreover, the internal and external iliac arteries were connected aberrantly to the dorsal aorta (Figure 6C), suggesting that the common iliac artery was also missing in the mutants. Together, these findings demonstrate that *Wnt2* plays a previously unknown role in regulating development of both the bladder and the umbilical arteries.

**Figure 6.**
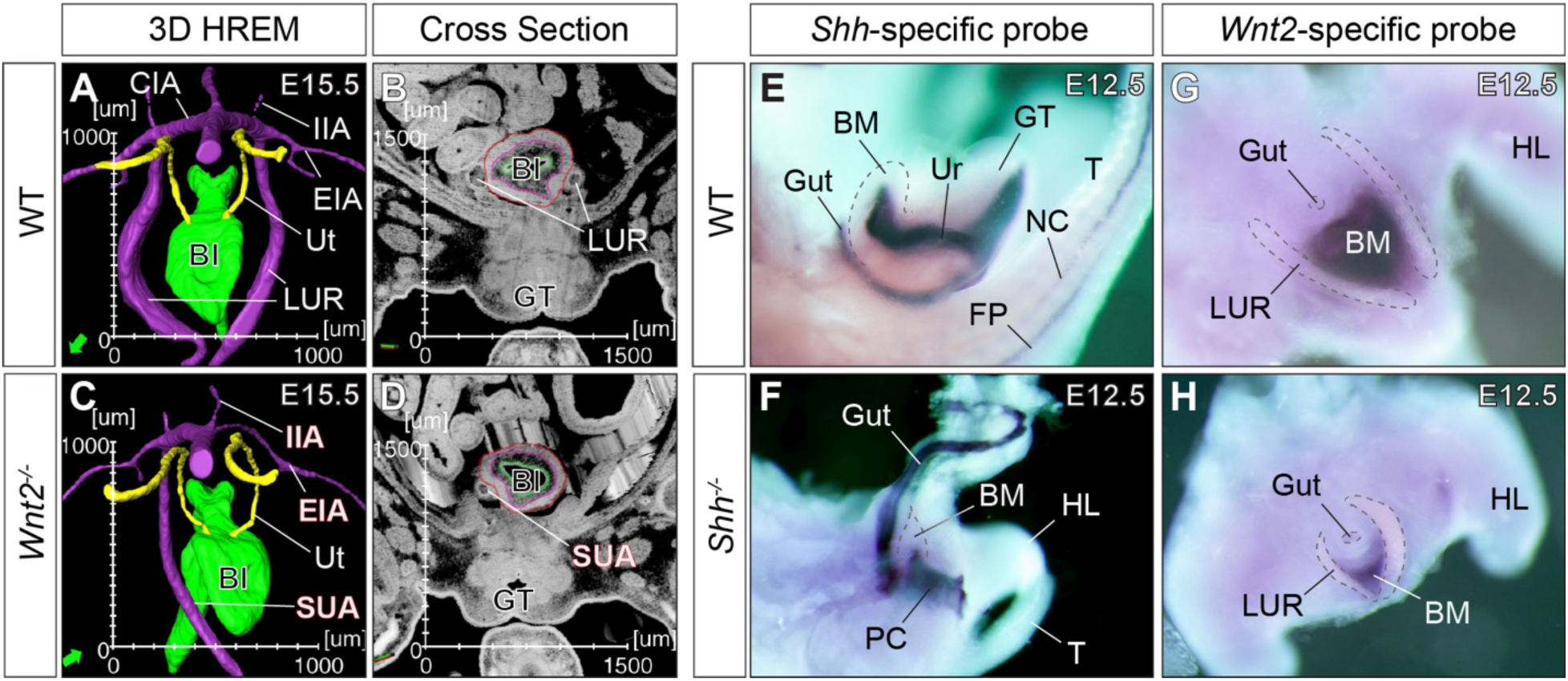
The *Shh-Wnt2* molecular pathway orchestrates co-development of the bladder and umbilical arteries. **A**-**D**, HREM visualizations reveal the single umbilical artery (SUA) and the iliac artery defects associated with *Wnt2^-/-^* mutants (C and D). **E**-**H**, *Wnt2* expression is reduced in *Shh^-/-^* mutants. Wholemount RNA *in situ* hybridization using gene-specific probes.

### Epithelial *Shh* regulates *Wnt2* expression in the bladder mesenchyme

We next examined the possibility whether *Shh* may regulate *Wnt2* gene expression during bladder development. Strong *Shh* expression was detected in the cloacal epithelium as well as the gut epithelium of wild type embryos (Figure 6E). The first axon of *Shh* gene is deleted in *Shh^-/-^* mutants, resulting in a null allele that produces truncated transcripts without any functional protein (39). High levels of truncated *Shh* transcripts were detected in *Shh^-/-^* mutant embryos (Figure 6F), indicating that *Shh* expression is independent to *Shh* signaling *in vivo*. Expression pattern of the truncated *Shh* transcripts showed that the rostral and caudal outgrowth was absent in *Shh^-/-^* mutant embryos (Figure 6F), consistent with HREM 3D findings. Strong *Wnt2* transcript signal was observed at the triangular mesenchymal space flanked by the umbilical arteries in wild type controls. Moreover, *Wnt2* signal was obviously weaker in *Shh^-/-^* embryos (Figures 6G vs H, rostral views), suggesting that epithelial *Shh* signal is required for *Wnt2* expression in the rostral outgrowth mesenchyme during bladder development.

## Discussion

In this study, we report that mammalian urinary bladder emerges as the rostral outgrowth of the cloaca (Figure 7). The outgrowth is mediated in part by the epithelial *Shh* and mesenchymal *Wnt2* signals. Unexpectedly, the outgrowth is required for formation of both the bladder and the umbilical arteries. In addition to supporting life after birth, our findings suggest that having the bladder may also provide survival advantage before birth by supporting development of the umbilical arteries.

**Figure 7.**
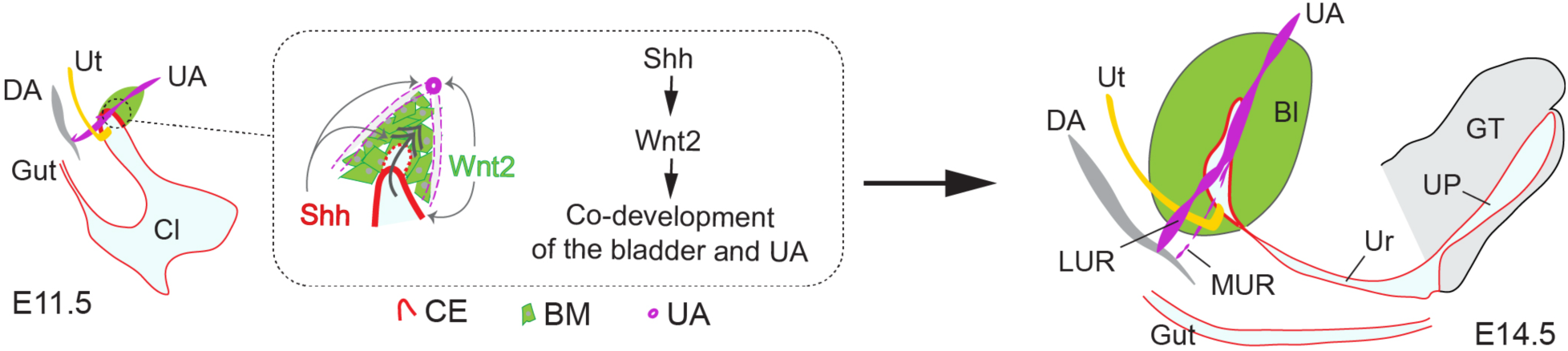
A proposed model of co-development of the bladder and umbilical arteries. Rostral outgrowth (double arrow) of the cloaca into the *Wnt2-*positive mesenchymal progenitors (green) results in formation of the bladder. A caudal outgrowth of the cloacal forms the urethra (U) and urethral plate (UP) of the genital tubercle (GT). Remodeling of the umbilical artery (UA) includes degeneration of the medial umbilical artery roots (MUR) and establishment of the lateral umbilical roots (LUR). *Shh* (red) from cloacal epithelium, which is required for *Wnt2* expression, controls both the bladder development and umbilical artery remodeling. *Wnt2* regulates proliferation of bladder mesenchyme and epithelium, as well as remodeling of the umbilical arteries. Together, the *Shh-Wnt2* pathway orchestrates co-development of the bladder and umbilical arteries. Feedback signal(s) from the umbilical arteries (purple) is expected to regulate bladder formation. See other figures for abbreviations.

The bladder appears at the junction between embryonic cloaca and extraembryonic allantois. While we could not rule out a possible contribution of the allantois, especially in humans (2, 26), findings here suggest the bladder originate from the rostral outgrowth of the cloaca in both humans and mice. For reptiles and birds, the allantoic gut is a vital extraembryonic organ that stores metabolic wastes before hatching. The allantois is conserved in mammals; however, human allantoic gut is underdeveloped and degenerated during fetal development. Our 3D morphometric and serial histological analyses have not identified any sign of the allantoic gut throughout mouse embryonic stages. The lack of allantoic gut strongly suggest that the bladder is less likely originated from the extraembryonic allantois in mice.

We show that the outgrowth extends into the triangular area located rostral to the cloaca. This previously uncharacterized triangular mesenchymal domain is flanked by the umbilical arteries and marked by *Wnt2* gene expression. The rostral outgrowth expands and becomes the bladder in mice (∼E13.5) and humans (∼CS18). Concomitant with bladder formation, we show that the umbilical arteries undergo dramatic remodeling: the MUR degenerates while the LUR emerges. Remodeling of the umbilical arteries occurs in both human and mouse and takes place at the comparable embryonic stages, implying a conserved mechanism underlying the co-development. This paradigm implies that signal(s) required for bladder development may also regulate formation of the umbilical arteries. The umbilical arteries may also direct development of the bladder.

The proposed moonlighting role of embryonic bladder predicts that lack of bladder is incompatible to life. The onset of bladder development and umbilical artery remodeling takes place during the second month of gestation in humans. It is worth noting that nearly 80% of miscarriages in human pregnancies occurs in the first trimester with unknown etiology (44-46). An impaired co-development program might be an overlooked reason for those miscarriages. Future studies are needed to determine to what extent the molecular network underlying normal bladder development may contribute to development of the umbilical arteries and, ultimately, survival of mammalian conceptus and successful pregnancies.

## Acknowledgments

We thank past and current members of the X.L. lab at Boston Children’s Hospital/Harvard Medical School at Boston and Cedars-Sinai Medical Center at Los Angeles, particularly Drs. Chunming Guo, Yichen Huang, and Satoshi Kaneko for technical supports and helpful discussions. We are grateful for Christa Lam for critical and insightful comments on the manuscript. This work was supported by NIH/NIDDK (R01DK110477 and U01DK131377, X.L.), NIH/NCI (1R01CA267108 and 1P01CA278732, X.L.), and NHLBI (1R01HL136921, X.L.).

## Materials and Methods

### Mice

All animal studies were reviewed and approved by the Institutional Animal Care and Use Committee at Cedars-Sinai Medical Center. Wild type C57BL/6 mice were obtained from Charles River Laboratories. *Shh^+/-^*(B6.Cg-*Shh^tm1(EGFP/cre)Cjt/J^*, 005622) and *R26R^lacZ^*B6.129S4-*Gt(ROSA)26^Sortm1Sor/J^*, 003474) were obtained from the Jackson Laboratory. The *Wnt2* mouse line, as previously described (41), was generously provided by Dr. Edward Morrisey from the University of Pennsylvania.

### High-resolution episcopic microscopy (HREM) and 3D reconstruction

HREM was performed as previously described (6). Briefly, staged mouse embryos (wild type, *Shh^-/-^*, and *Wnt2^-/-^* mutants) were fixed in 10% buffer formalin (>1 day) and embedded in JB-4 embedding Kit (Polyscience) containing eosin (Fisher Scientific) and acridine orange (Sigma Aldrich). High-resolution serial block face images were collected using a stereo zoom microscope (Zeiss, Axio Zoom.V16) using a Hamamatsu digital camera (Photonics K.K., C11440-10C) from both green and red fluorescent channels. All images were inverted using Photoshop (Adobe, V13.0) and further converted to a volume data using the 3-D visualization software (Amira V5.4.5) to directly visualize size and morphometric features of each specimen. Amira files of digitized human embryos from Carnegie stages (CS) 8 to 21, as reported previously (8), were downloaded directly from http://3datlasofhumanembryology.com. Structures, including the cloaca, bladder, and the umbilical arteries were manually annotated. The resulting structural segmentations were used in quantitative measurements including distance, surface area, and volume. Animations were converted into movie files using the MovieMaker module.

### Histology, immunostaining, and wholemount RNA *in situ* hybridization

The same procedures that we described previously were used (24, 47). Briefly, mouse embryos were micro-dissected in cold phosphate buffer saline and fixed in 10% buffer formalin for the subsequent analyses. Hematoxylin and eosin (H&E) staining was performed on serial histological sections of E8.5 embryos. Midline sagittal sections of E15.5 wildtype and *Wnt2^-/-^* mutants were stained using phospho-histone H3 (pHH3, Upstate, 06-570) and smooth muscle actin (SMA, Sigma, A2547)-specific antibodies. Wholemount RNA *in situ* hybridization of staged embryos was done using *Wnt2* and *Shh-* specific riboprobes as previously described (41, 47). Expression patterns of the representative signature genes were manually curated using the publicly available GenePaint database (https://gp3.mpg.de)(43).

---

**Table S1, related to Figure 1.** List of HREM digitized human embryos.

| CS | Specimen# | Origin | Year | CRL (mm) | Day | Sex | Plane | Sections# | Pixel size(μm) | Z-res (μm) |
| --- | --- | --- | --- | --- | --- | --- | --- | --- | --- | --- |
| 8 | 8671 | CC | 1949 | 0.61 |  |  | T | 226 | 0.86 | 2.70 |
| 9 | H712 | BC | 1957 | 1.57 |  |  | T | 437 | 1.81 | 3.59 |
| 10 | 6330 | CC | 1931 | 1.95 | 28 |  | T | 168 | 0.54 | 11.63 |
| 11 | 6784 | CC | 1933 | 2.46 |  |  | T | 283 | 0.86 | 8.70 |
| 12 | 8505A | CC | 1947 | 2.86 |  |  | T | 277 | 1.08 | 10.32 |
| 13 | 836 | CC | 1914 | 4.09 | 32 | F | T | 247 | 5.89 | 16.55 |
| 14 | 8314 | CC | 1945 | 5.16 | 22 |  | T | 639 | 2.23 | 8.07 |
| 15 | 3512 | CC | 1921 | 6.55 |  |  | T | 651 | 2.23 | 10.06 |
| 16 | 6517 | CC | 1931 | 10.46 | 39 |  | T | 547 | 5.40 | 19.13 |
| 17 | 6521 | CC | 1933 | 10.60 |  |  | T | 1059 | 4.48 | 10.01 |
| 18 | 6524 | CC | 1933 | 9.73 |  |  | T | 956 | 4.48 | 10.18 |
| 20 | 462 | CC | 1910 | 15.93 |  | M | T | 376 | 4.48 | 42.36 |
| 21 | 7254 | CC | 1936 | 17.36 |  | M | T | 288 | 4.48 | 60.12 |

CS: Carnegie stage; Origin of the specimen: CC: Carnegie Collection, BC: Boyd Collection; Year: Year of acquisition; CRL: Crown-rump-length in mm; Day: days post ovulation; Plane of sectioning: O = Oblique, T= Transversal; Z-res: Calculated Z-resolution in μm.
de Bakker BS, de Jong KH, Hagoort J, de Bree K, Besselink CT, de Kanter FE, Veldhuis T, Bais B, Schildmeijer R, Ruijter JM, Oostra RJ, Christoffels VM, Mooman AF. An interactive three-dimensional digital atlas and quantitative database of human development. Science. 2016 Nov 25; 354 (6315): aag0053.

**Table S2, related to Figure 1.** List of HREM digitized mouse embryos.

| **Specimen#** | **Embryonic days** | **Somite#** | **Sections#** | **Pixel size(μm)** | **Z-res (μm)** |
| --- | --- | --- | --- | --- | --- |
| 5-6 | 8.5(2X with YS) |  | 3810 | 3.63 | 2.00 |
| 122 | 9.5a | 27 | 1567 | 1.80 | 1.50 |
| 123 | 9.5b | 27 | 1492 | 1.50 | 1.50 |
| 124 | 9.5c | 27 | 1446 | 1.60 | 1.50 |
| 1908 | 10.0a | 31 | 1740 | 1.50 | 1.50 |
| 145 | 10.0b | 30 | 1742 | 1.70 | 1.50 |
| 147 | 10.0c | 32 | 1811 | 1.60 | 1.50 |
| 78 | 10.5a | 36 | 1995 | 2.10 | 2.00 |
| 42 | 10.5b | 35 | 1458 | 2.10 | 2.10 |
| 37 | 10.5c | 37 | 1650 | 2.00 | 2.00 |
| 57 | 11.0a | 41 | 2179 | 2.40 | 2.00 |
| 58 | 11.0b | 43 | 2374 | 2.50 | 2.00 |
| 56 | 11.5a | 46 | 2477 | 3.00 | 2.00 |
| 83 | 11.5b | 45 | 2349 | 2.30 | 2.00 |
| 66 | 11.5c | 46 | 2682 | 2.80 | 2.00 |
| 125 | 12.0a | 48 | 2602 | 2.80 | 2.00 |
| 86 | 12.0b | 48 | 2578 | 2.60 | 2.00 |
| 126 | 12.0c | 48 | 2776 | 2.40 | 2.00 |
| 34 | 12.5a | >50 | 2533 | 3.20 | 2.50 |
| 89 | 12.5b | >50 | 2552 | 2.90 | 2.50 |
| 128 | 12.5c | >50 | 2797 | 2.90 | 2.50 |
| 141 | 13.0a | >55 | 3247 | 3.20 | 2.50 |
| 142 | 13.0b | >55 | 3237 | 3.50 | 2.50 |
| 157 | 13.0c | >55 | 3511 | 3.20 | 2.50 |
| 72 | 13.5a | >55 | 3006 | 3.80 | 2.50 |
| 73 | 13.5b | >55 | 2997 | 3.60 | 2.50 |
| 77 | 13.5c | >55 | 3359 | 3.80 | 2.50 |

**Figure S1, Related to Figure 1.**
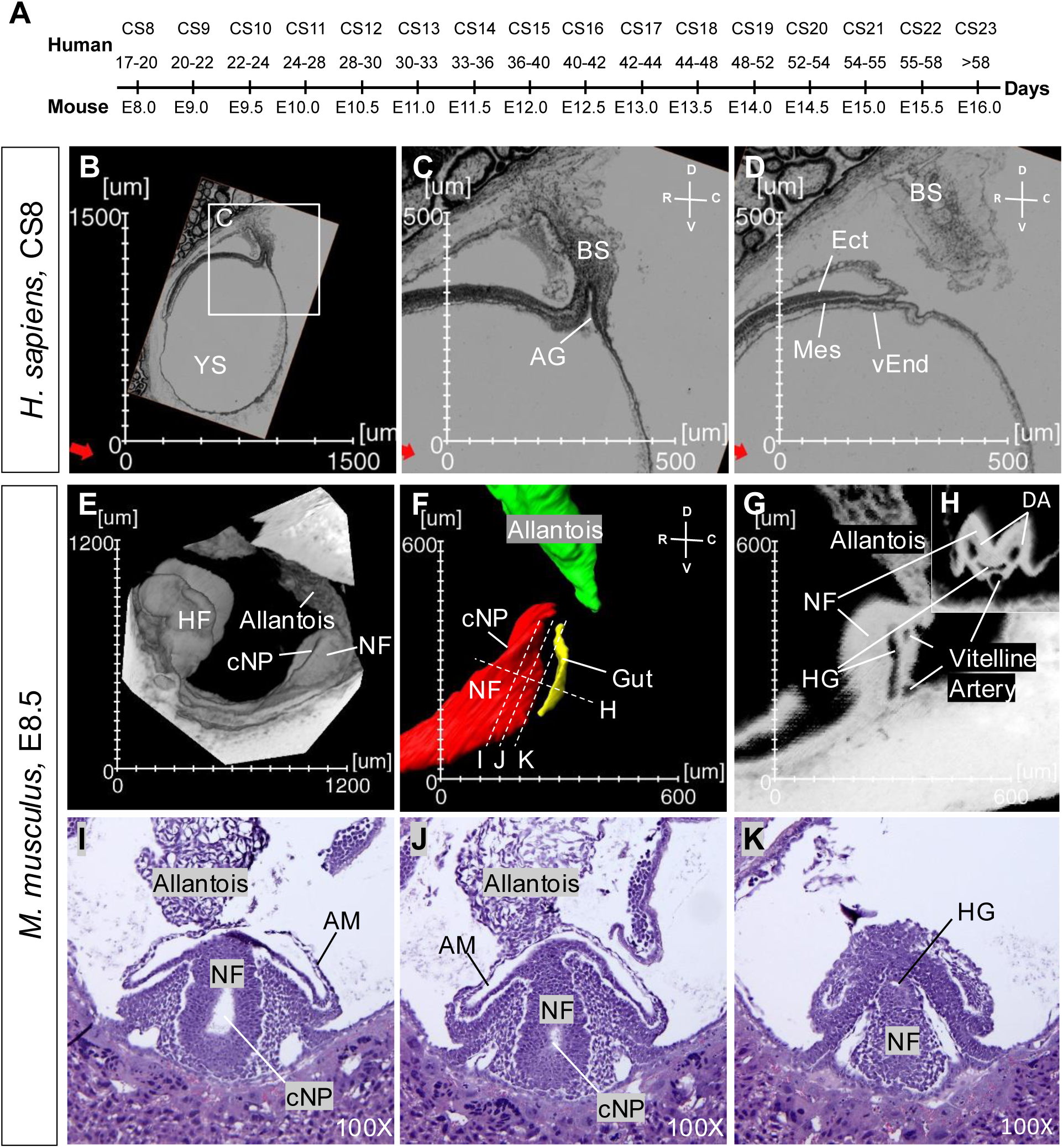
Mouse embryos absent of the allantoic gut. **A**, Comparative developmental stages of human and mouse embryos. CS, Carnegie stage; E, embryonic day. **B-D**, Digital sagittal sections of CS8 human embryo through the allantois (B and C) and at the lateral side (D). **E-G**, Digital sagittal views of wholemount (E), 3D segmentation (F), and midline section (G) of E8.5 mouse embryo. **H**, Digital cross section of E8.5 embryos from the plane shown in F. **I-K**, Serial histologically stained sections of E8.5 mouse embryos. Locations and orientations of the sections are shown in F. AM, amnion; BS, body stalk; cNP, caudal neural pore; Ect, ectoderm; HF, headfold; Mes, mesoderm; vEnd, visceral endoderm; NF, neural fold; YS, yolk sac. See Figure 1 for additional abbreviations.

**Figure S2,** Video related to Figure 1. 3D visualization of the developing urinary tract and the associated arterial network of human embryos at CS12, 14, 18, and 21.

**Figure S3,** Video related to Figures 1-3. 3D visualization of the developing mouse cloaca and the associated arterial network from E10.5, 11.5, 13.5, to E15.5.

**Figure S4,** Video related to Figure 4. 3D visualization of the developmental defects of the bladder and umbilical arteries of *Shh^-/-^*mutant mouse embryos at E10.5, 11.5, 12.5, and E13.5. *, indicating absence of the bladder.

**Figure S5,** Video related to Figure 4. 3D visualization of the *Shh-*positive (*Shh^+^*) lineages at E11.5. Double heterozygous *Shh^+/-^;R26R^lacZ^* embryos were stained with X-gal to detect the Shh^+^ lineages. Stained cells were visualized directly using HREM to create the 3D views.

**Figure S6, Related to Figure 5.**
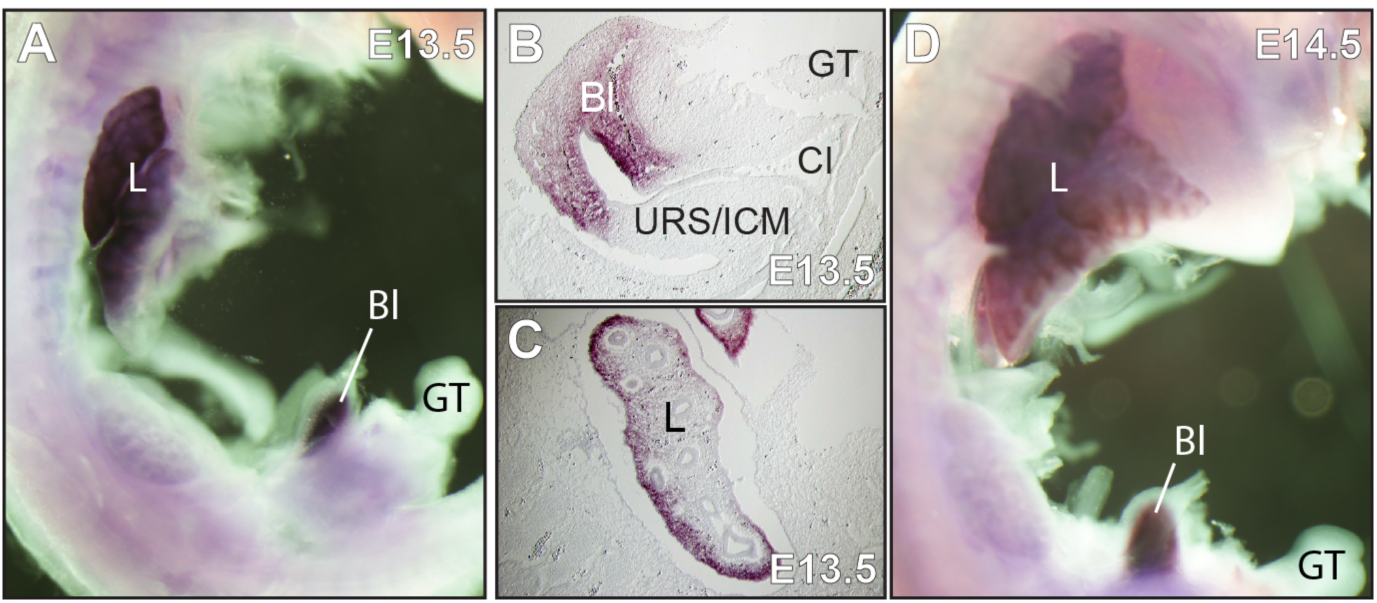
*Wnt2* is specifically expressed in mesenchymal progenitors of the developing bladder and lung. Wholemount RNA *in situ* hybridization using a *Wnt2-*specific probe (dark purple staining). **A-C**, A wholemount (A) and cryostat section (B and C) views of E13.5 mouse embryo after *Wnt2 in situ* hybridization. **D**, A wholemount view of E14.5 embryos after *Wnt2 in situ* hybridization.

## References

1. Hicks RM. The mammalian urinary bladder: an accommodating organ. Biological reviews of the Cambridge Philosophical Society. 1975;50(2):215–46.

2. Bentley PJ. The vertebrate urinary bladder: osmoregulatory and other uses. Yale J Biol Med. 1979;52(6):563–8.

3. McCarthy M, and McCarthy L. The evolution of the urinary bladder as a storage organ: scent trails and selective pressure of the first land animals in a computational simulation. SN Applied Sciences. 2019;1(12):1727.

4. Rasouly HM, and Lu W. Lower urinary tract development and disease. Wiley Interdiscip Rev Syst Biol Med. 2013;5(3):307–42.

5. Georgas KM, Armstrong J, Keast JR, Larkins CE, McHugh KM, Southard-Smith EM, et al. An illustrated anatomical ontology of the developing mouse lower urogenital tract. Development. 2015;142(10):1893–908.

6. Huang YC, Chen F, and Li X. Clarification of mammalian cloacal morphogenesis using high-resolution episcopic microscopy. Dev Biol. 2016;409(1):106–13.

7. Kruepunga N, Hikspoors J, Mekonen HK, Mommen GMC, Meemon K, Weerachatyanukul W, et al. The development of the cloaca in the human embryo. J Anat. 2018;233(6):724–39.

8. de Bakker BS, de Jong KH, Hagoort J, de Bree K, Besselink CT, de Kanter FE, et al. An interactive three-dimensional digital atlas and quantitative database of human development. Science. 2016;354(6315).

9. Glenn JF. Agenesis of the bladder. J Am Med Assoc. 1959;169(17):2016–8.

10. Yahya MH. Bladder Agenesis: A Systematic Review. Cureus. 2023;15(9):e45121.

11. Dykes EH, Oesch I, Ransley PG, and Hendren WH. Abnormal aorta and iliac arteries in children with urogenital abnormalities. J Pediatr Surg. 1993;28(5):696–700.

12. Lowrey T, Josephs S, and Baker LA. Bladder Agenesis and Associated Pelvic Arterial Anomaly in 2 Female Pediatric Patients. Urology. 2019;123:227–9.

13. Thottungal AD, Charles AK, Dickinson JE, and Bower C. Caudal dysgenesis and sirenomelia-single centre experience suggests common pathogenic basis. Am J Med Genet A. 2010;152A(10):2578-87.

14. Garrido-Allepuz C, Haro E, Gonzalez-Lamuno D, Martinez-Frias ML, Bertocchini F, and Ros MA. A clinical and experimental overview of sirenomelia: insight into the mechanisms of congenital limb malformations. Disease models & mechanisms. 2011;4(3):289–99.

15. Tschopp P, Sherratt E, Sanger TJ, Groner AC, Aspiras AC, Hu JK, et al. A relative shift in cloacal location repositions external genitalia in amniote evolution. Nature. 2014;516(7531):391–4.

16. Herrera AM, and Cohn MJ. Embryonic origin and compartmental organization of the external genitalia. Scientific reports. 2014;4:6896.

17. Lozovska A, Korovesi AG, Dias A, Lopes A, Fowler DA, Martins GG, et al. Tgfbr1 controls developmental plasticity between the hindlimb and external genitalia by remodeling their regulatory landscape. Nat Commun. 2024;15(1):2509.

18. Thompson DJ, Molello JA, Strebing RJ, and Dyke IL. Teratogenicity of adriamycin and daunomycin in the rat and rabbit. Teratology. 1978;17(2):151–7.

19. Liu MI, Hutson JM, and Zhou B. Critical timing of bladder embryogenesis in an adriamycin-exposed rat fetal model: a clue to the origin of the bladder. J Pediatr Surg. 1999;34(11):1647–51.

20. Matsumaru D, Murashima A, Fukushima J, Senda S, Matsushita S, Nakagata N, et al. Systematic stereoscopic analyses for cloacal development: The origin of anorectal malformations. Scientific reports. 2015;5:13943.

21. Liaw A, Cunha GR, Shen J, Cao M, Liu G, Sinclair A, et al. Development of the human bladder and ureterovesical junction. Differentiation. 2018;103:66–73.

22. Wang C, Gargollo P, Guo C, Tang T, Mingin G, Sun Y, et al. Six1 and Eya1 are critical regulators of peri-cloacal mesenchymal progenitors during genitourinary tract development. Dev Biol. 2011;360(1):186–94.

23. Wang C, Wang J, Borer JG, and Li X. Embryonic origin and remodeling of the urinary and digestive outlets. PLoS One. 2013;8(2):e55587.

24. Guo C, Sun Y, Guo C, MacDonald BT, Borer JG, and Li X. Dkk1 in the peri-cloaca mesenchyme regulates formation of anorectal and genitourinary tracts. Dev Biol. 2014;385(1):41–51.

25. 25. Mossman HW. Vertebrate Fetal Membranes. Rutgers University Press, New Brunswick, New Jersey; 1987.

26. van der Putte SC. Anal and ano-urogenital malformations: a histopathological study of "imperforate anus" with a reconstruction of the pathogenesis. Pediatr Dev Pathol. 2006;9(4):280–96.

27. Zeng H, and Liu A. TMEM132A regulates mouse hindgut morphogenesis and caudal development. Development. 2023;150(14).

28. Haraguchi R, Mo R, Hui C, Motoyama J, Makino S, Shiroishi T, et al. Unique functions of Sonic hedgehog signaling during external genitalia development. Development. 2001;128(21):4241–50.

29. Perriton CL, Powles N, Chiang C, Maconochie MK, and Cohn MJ. Sonic hedgehog signaling from the urethral epithelium controls external genital development. Dev Biol. 2002;247(1):26–46.

30. Haraguchi R, Motoyama J, Sasaki H, Satoh Y, Miyagawa S, Nakagata N, et al. Molecular analysis of coordinated bladder and urogenital organ formation by Hedgehog signaling. Development. 2007;134(3):525–33.

31. Garrido-Allepuz C, Gonzalez-Lamuno D, and Ros MA. Sirenomelia phenotype in bmp7;shh compound mutants: a novel experimental model for studies of caudal body malformations. PLoS One. 2012;7(9):e44962.

32. Zakin L, Reversade B, Kuroda H, Lyons KM, and De Robertis EM. Sirenomelia in Bmp7 and Tsg compound mutant mice: requirement for Bmp signaling in the development of ventral posterior mesoderm. Development. 2005;132(10):2489–99.

33. Suzuki K, Adachi Y, Numata T, Nakada S, Yanagita M, Nakagata N, et al. Reduced BMP signaling results in hindlimb fusion with lethal pelvic/urogenital organ aplasia: a new mouse model of sirenomelia. PLoS One. 2012;7(9):e43453.

34. Hikspoors JP, Soffers JH, Mekonen HK, Cornillie P, Kohler SE, and Lamers WH. Development of the human infrahepatic inferior caval and azygos venous systems. J Anat. 2015;226(2):113–25.

35. Seifert AW, Harfe BD, and Cohn MJ. Cell lineage analysis demonstrates an endodermal origin of the distal urethra and perineum. Dev Biol. 2008;318(1):143–52.

36. Downs KM, and Rodriguez AM. The mouse fetal-placental arterial connection: A paradigm involving the primitive streak and visceral endoderm with implications for human development. Wiley interdisciplinary reviews Developmental biology. 2020;9(2):e362.

37. Chapouly C, Guimbal S, Hollier PL, and Renault MA. Role of Hedgehog Signaling in Vasculature Development, Differentiation, and Maintenance. Int J Mol Sci. 2019;20(12).

38. Kolesova H, Roelink H, and Grim M. Sonic hedgehog is required for the assembly and remodeling of branchial arch blood vessels. Dev Dyn. 2008;237(7):1923–34.

39. Harfe BD, Scherz PJ, Nissim S, Tian H, McMahon AP, and Tabin CJ. Evidence for an expansion-based temporal Shh gradient in specifying vertebrate digit identities. Cell. 2004;118(4):517–28.

40. Jasmine, Baraiya DH, Kavya TT, Mandal A, Chakraborty S, Sathish N, et al. Epithelial and mesenchymal compartments of the developing bladder and urethra display spatially distinct gene expression patterns. Dev Biol. 2025;520:155–70.

41. Goss AM, Tian Y, Tsukiyama T, Cohen ED, Zhou D, Lu MM, et al. Wnt2/2b and beta-catenin signaling are necessary and sufficient to specify lung progenitors in the foregut. Dev Cell. 2009;17(2):290–8.

42. Peng T, Tian Y, Boogerd CJ, Lu MM, Kadzik RS, Stewart KM, et al. Coordination of heart and lung co-development by a multipotent cardiopulmonary progenitor. Nature. 2013;500(7464):589–92.

43. Visel A, Thaller C, and Eichele G. GenePaint.org: an atlas of gene expression patterns in the mouse embryo. Nucleic Acids Res. 2004;32(Database issue):D552-6.

44. Wilcox AJ, Weinberg CR, O’Connor JF, Baird DD, Schlatterer JP, Canfield RE, et al. Incidence of early loss of pregnancy. The New England journal of medicine. 1988;319(4):189–94.

45. Ammon Avalos L, Galindo C, and Li DK. A systematic review to calculate background miscarriage rates using life table analysis. *Birth defects research Part A*, Clinical and molecular teratology. 2012;94(6):417–23.

46. Mukherjee S, Velez Edwards DR, Baird DD, Savitz DA, and Hartmann KE. Risk of miscarriage among black women and white women in a U.S. Prospective Cohort Study. Am J Epidemiol. 2013;177(11):1271–8.

47. Guo C, Balsara ZR, Hill WG, and Li X. Stage- and subunit-specific functions of polycomb repressive complex 2 in bladder urothelial formation and regeneration. Development. 2017;144(3):400–8.

